# Hybrid genome assembly and annotation of *Danionella translucida*

**DOI:** 10.1101/539692

**Authors:** Mykola Kadobianskyi, Lisanne Schulze, Markus Schuelke, Benjamin Judkewitz

**Affiliations:** Einstein Center for Neurosciences, NeuroCure Cluster of Excellence, Charité – Universitätsmedizin Berlin, Charitéplatz 1, 10117 Berlin, Germany.

**Keywords:** Hybrid genome assembly, Danionella translucida

## Abstract

Studying neuronal circuits at cellular resolution is very challenging in vertebrates due to the size and optical turbidity of their brains. *Danionella translucida*, a close relative of zebrafish, was recently introduced as a model organism for investigating neural network interactions in adult individuals. *Danionella* remains transparent throughout its life, has the smallest known vertebrate brain and possesses a rich repertoire of complex behaviours. Here we sequenced, assembled and annotated the *Danionella translucida* genome employing a hybrid Illumina/ Nanopore read library as well as RNA-seq of embryonic, larval and adult mRNA. We achieved high assembly continuity using low-coverage long-read data and annotated a large fraction of the transcriptome. This dataset will pave the way for molecular research and targeted genetic manipulation of this novel model organism.

## Introduction

The size and opacity of vertebrate tissues limit optical access to the brain and hinder investigations of intact neuronal networks *in vivo*. As a result, many scientists focus on small, superficial brain areas, such as parts of the cerebral cortex in rodents, or on early developmental stages of small transparent organisms, like zebrafish larvae. In order to overcome these limitations, we recently developed as a novel model organism for the optical investigation of neuronal circuit activity in vertebrates – *Danionella translucida* (DT), a transparent cyprinid fish (1, 2) with the smallest known vertebrate brain (3, 4). The majority of DT tissues remain transparent throughout its life (Fig. 1). DT displays a rich repertoire of social behaviours, such as schooling and vocal communication, and is amenable to genetic manipulation using genetic tools that are already established in zebrafish. As such, this species is a promising model organism for studying the function of neuronal circuits across the entire brain. Yet, a continuous annotated genome reference is still needed to enable targeted genetic and transgenic studies and facilitate the adoption of DT as a model organism.

**Fig. 1.**
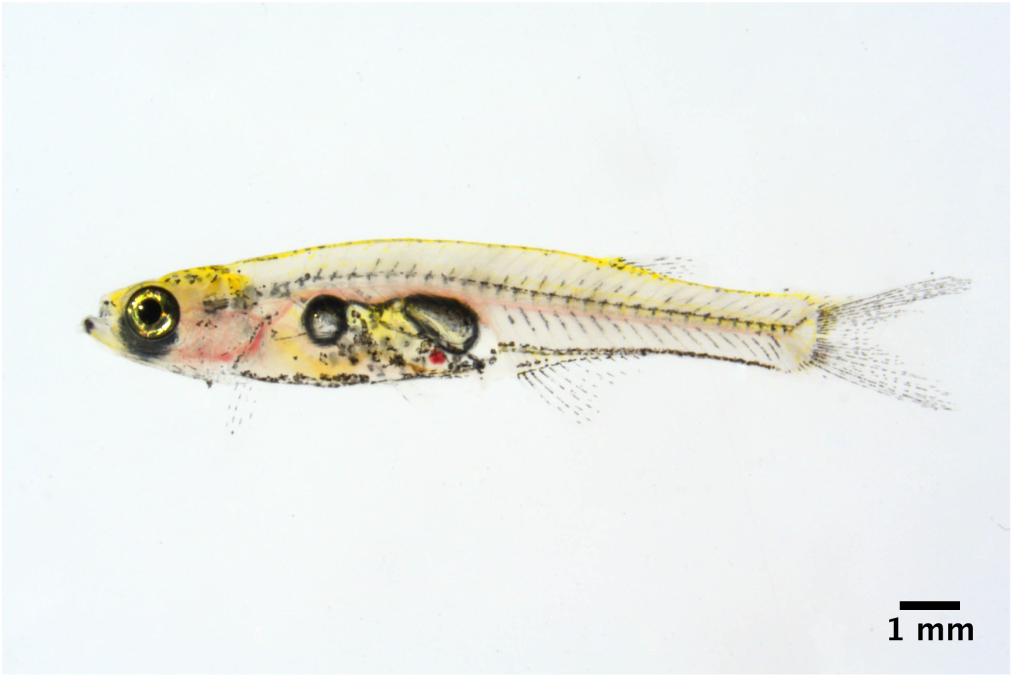
Male adult *Danionella translucida* showing transparency.

Next-generation short-read sequencing advances steadily decreased the price of the whole-genome sequencing and enabled a variety of genomic and metagenomic studies. However, short-read-only assemblies often struggle with repetitive and intergenic regions, resulting in fragmented assembly and poor access to regulatory and promoter sequences (5, 6). Long-read techniques, such as PacBio and Nanopore, can generate reads up to 2 Mb (7), but they are prone to errors, including frequent indels, which can lead to artefacts in long-read-only assemblies (6). Combining short- and long-read sequencing technologies in hybrid assemblies recently produced high-quality genomes in fish (8, 9).

Here we report the hybrid Illumina/Nanopore-based assembly of the *Danionella translucida* genome. A combination of deep-coverage Illumina sequencing with a single Nanopore sequencing run produced an assembly with scaffold N50 of 340 kb and Benchmarking Universal Single-Copy Orthologs (BUSCO) genome completeness score of 92%. Short- and long-read RNA sequencing data used together with other fish species annotated proteomes produced an annotation dataset with BUSCO transcriptome completeness score of 86%.

### Genomic sequencing libraries

For genomic DNA sequencing we generated paired-end and mate-pair Illumina sequencing libraries and one Nanopore library. We extracted DNA from fresh DT tissues with phenolchloroform-isoamyl alcohol. For Illumina sequencing, we used 5 days post fertilisation (dpf) old larvae. A shotgun paired-end library with ~500 bp insert size was prepared with TruSeq kit (Illumina). Sequencing on HiSeq 4000 generated 1.347 billion paired-end reads. A long 10 kb mate-pair library was prepared using the Nextera Mate Pair Sample Prep Kit and sequenced on HiSeq 4000, resulting in 554 million paired-end reads. Read library quality was controlled using FastQC v0.11.8 (10).

A Nanopore sequencing high-molecular-weight gDNA library was prepared from 3 months post fertilisation (mpf) DT tails. We used ~400 ng of DNA with the 1D Rapid Sequencing Kit according to manufacturer’s instructions to produce the longest possible reads. This library was sequenced with the MinION sequencer on a single R9.4 flowcell using MinKNOW software for sequencing and base-calling, producing a total of 4.3 Gb sequence over 825k reads. The read library N50 was 11.6 kb with the longest read being ~200 kb. Sequencing data statistics are summarised in Table 1.

**Table 1.**
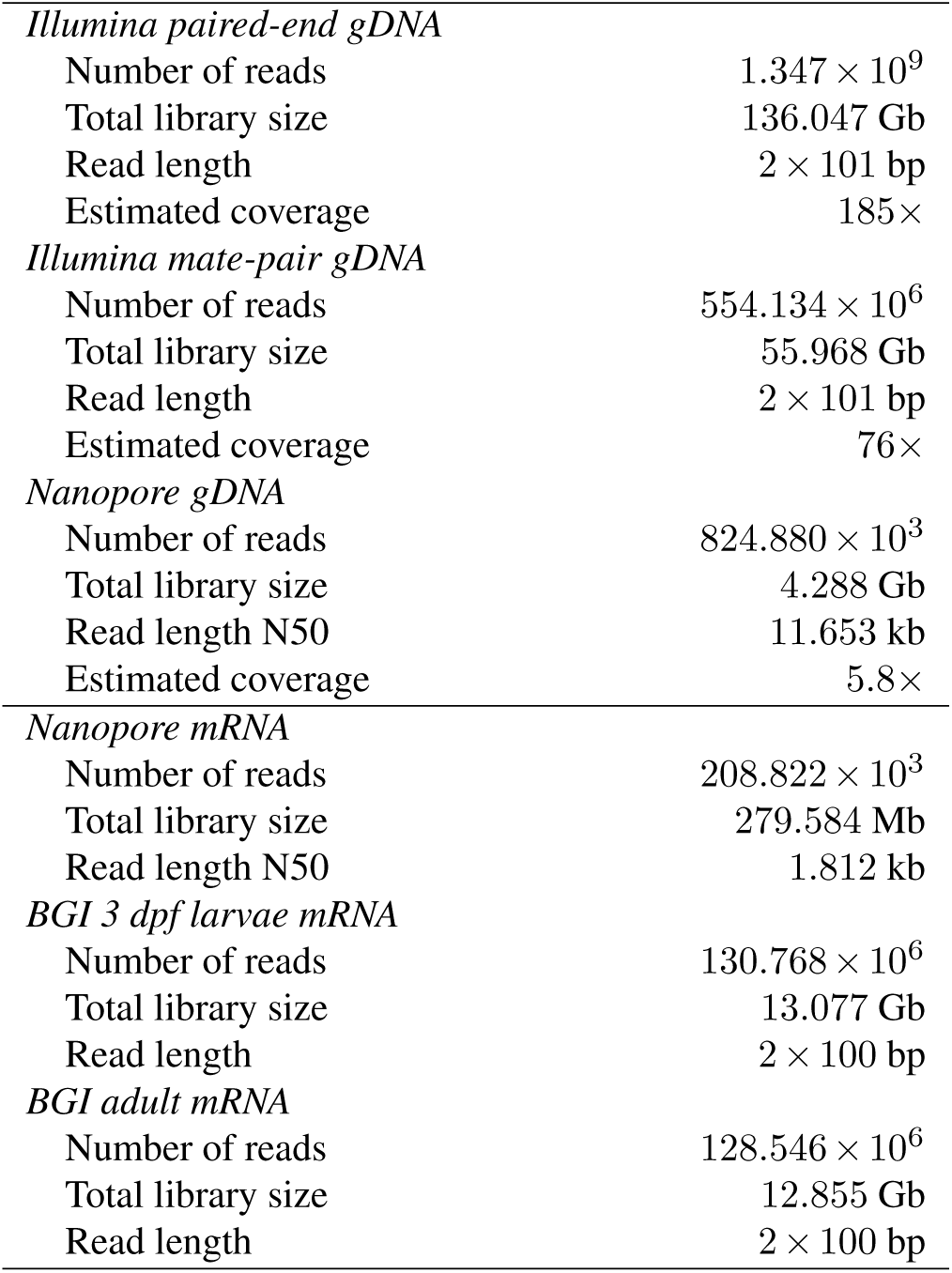
Sequencing library statistics. gDNA stands for genomic DNA sequencing, mRNA for poly-A tailed RNA sequencing.

### Genome assembly

The genome assembly and annotation pipeline is shown in Fig. 2. We estimated the genome size using the k-mer histogram method with Kmergenie v1.7016 on the paired-end Illumina library preprocessed with fast-mcf v1.04.807 (11, 12), which produced a putative assembly size of approximately 750 Mb. This translates into 185-fold Illumina and 5.8-fold Nanopore sequencing depths.

**Fig. 2.**
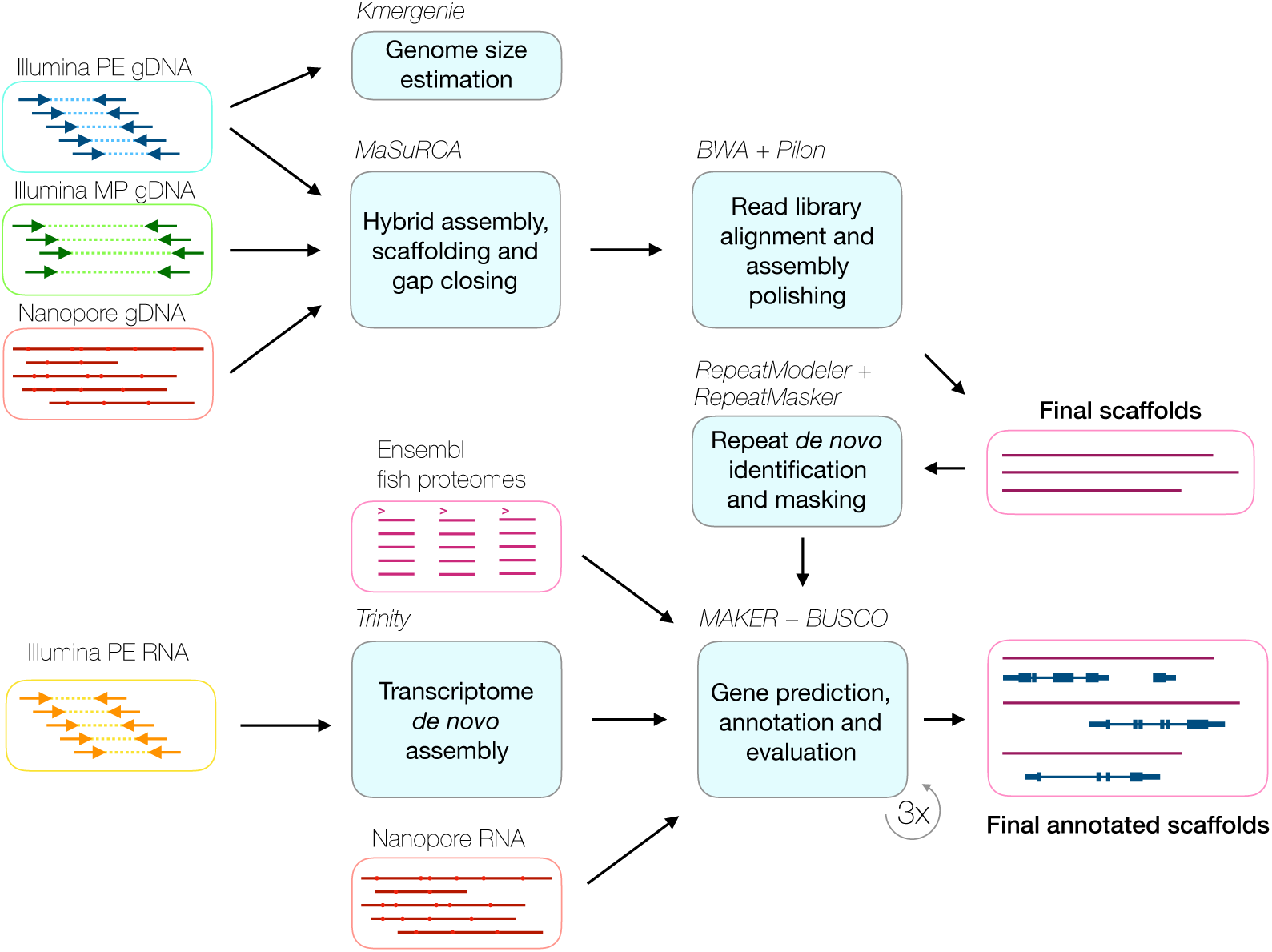
DT genome assembly and annotation pipeline. **PE**, paired-end; **MP**, mate-pair.

Multiple published assembly pipelines utilise a combination of short- and long-read sequencing. Our assembler of choice was MaSuRCA v3.2.6 (13), since it has already been used to generate high-quality assemblies of fish genomes, providing a large continuity boost even with low amount of input Nanopore reads (8, 9). Briefly, Illumina paired-end shotgun reads were non-ambiguously extended into the superreads, which were mapped to Nanopore reads for error correction, resulting in megareads. These megareads were then fed to the modified CABOG assembler that assembles them into contigs and, ultimately, mate-pair reads were used to do scaf- and gap repair.

Following MaSuRCA author’s recommendation (8), we have turned off the *frgcorr* module and provided raw read libraries for in-built read preprocessing. The initial genome assembly size estimated with the Jellyfish assembler module was 938 Mb. After the MaSuRCA pipeline processing we have polished the assembly with one round of Pilon v1.22, which attempts to resolve assembly errors and fill scaffold gaps using preprocessed reads mapped to the assembly (14). Statistics of the resulting assembly were generated using bbmap stats toolkit v37.32 (15) and are presented in Table 2.

**Table 2.** DT genome assembly statistics and completeness.

| Genome assembly statistics |  |
| --- | --- |
| Total scaffolds | 27,814 |
| Total contigs | 36,191 |
| Total scaffold sequence | 735.373 Mb |
| Total contig sequence | 725.755 Mb |
| Gap sequences | 1.308% |
| Scaffold N50 | 340.819 kb |
| Contig N50 | 133.131 kb |
| Longest scaffold | 3.085 Mb |
| Longest contig | 995.155 kb |
| Fraction of genome in > 50 kb scaffolds | 88.3% |
| BUSCO genome completeness score |  |
| Complete | 91.5% |
| Single | 87.0% |
| Duplicated | 4.5% |
| Fragmented | 3.6% |
| Missing | 4.9% |
| Total number of orthologs (Actinopterygii) | 4,584 |

The resulting 735 Mb assembly had a scaffold N50 of 341 kb, the longest scaffold being more than 3 Mb. To assess the completeness of the assembly we used BUSCO v3 (16) with the Actinopterygii ortholog dataset. In total, 91.5% of the orthologs were found in the assembly.

### Transcriptome sequencing and annotation

We used three sources of transcriptome evidence for the DT genome annotation: (i) assembled poly-A-tailed short-read and raw Nanopore RNA sequencing libraries, (ii) protein databases from sequenced and annotated fish species and (iii) trained gene prediction software. For Nanopore RNA sequencing we extracted total nucleic acids from 1-2 dpf embryos using phenol-chloroform-isoamyl alcohol extraction followed by DNA digestion with DNAse I. Resulting total RNA was converted to double-stranded cDNA using poly-A selection at the reverse transcription step with the Maxima H Minus Double-Stranded cDNA Synthesis Kit (ThermoFisher). The double-stranded cDNA sequencing library was prepared and sequenced in the same way as the genomic DNA, resulting in 190 Mb sequence data distributed over 209k reads. These reads were filtered to remove 10% of the shortest ones. For short-read RNA-sequencing, we have extracted total RNA with the TRIzol reagent (Invitrogen) from 3 dpf larvae and from adult fish. RNA was poly-A enriched and sequenced as 100 bp paired-end reads on the BGISEQ-500 platform. After preprocessing this resulted in 65.4 million read pairs for 3 dpf larvae and in 64.3 million read pairs for adult fish specimens (Table 1). We first assembled the 100 bp paired-end RNA-seq reads *de novo* using Trinity v2.8.4 assembler (17). Resulting RNA contigs, together with the Nanopore cDNA reads and proteomes of 11 fish species from Ensembl (18) were used as the transcript evidence in MAKER v2.31.10 annotation pipeline (19). Repetitive regions were masked using a *de novo* generated DT repeat library (RepeatModeler v1.0.11 (20)). The highest quality annotations with average annotation distance (AED) < 0.25 were used to train SNAP (21) and Augustus (22) gene predictors. Gene models were then polished over two additional rounds of re-training and re-annotation. The final set of annotations consisted of 24,099 gene models with an average length of 13.4 kb and an average AED of 0.18 (Table 3). We added putative protein functions using MAKER from the UniProt database (23) and protein domains from the interproscan v5.30-69.0 database (24). tRNAs were searched for and annotated using tRNAscan-SE v1.4 (25). The BUSCO transcriptome completeness search found 86% of complete *Actinopterygii* orthologs. An example Interactive Genomics Viewer (IGV) v2.4.3 (26) window with the *dnmt1* gene is shown on Fig. 3, demonstrating the annotation and RNA-seq coverage.

**Table 3.** DT transcriptome annotation statistics.

|  |  |
| --- | --- |
| Total gene models | 24,099 |
| Total functionally annotated gene models | 21,491 |
| Gene models with AED < 0.5 | 95% |
| Mean AED | 0.18 |
| BUSCO annotation completeness score |  |
| Complete | 86.3% |
| Single | 80.6% |
| Duplicated | 5.7% |
| Fragmented | 7.1% |
| Missing | 6.6% |
| Total number of orthologs (*Actinopterygii*) | 4,584 |

**Fig. 3.**
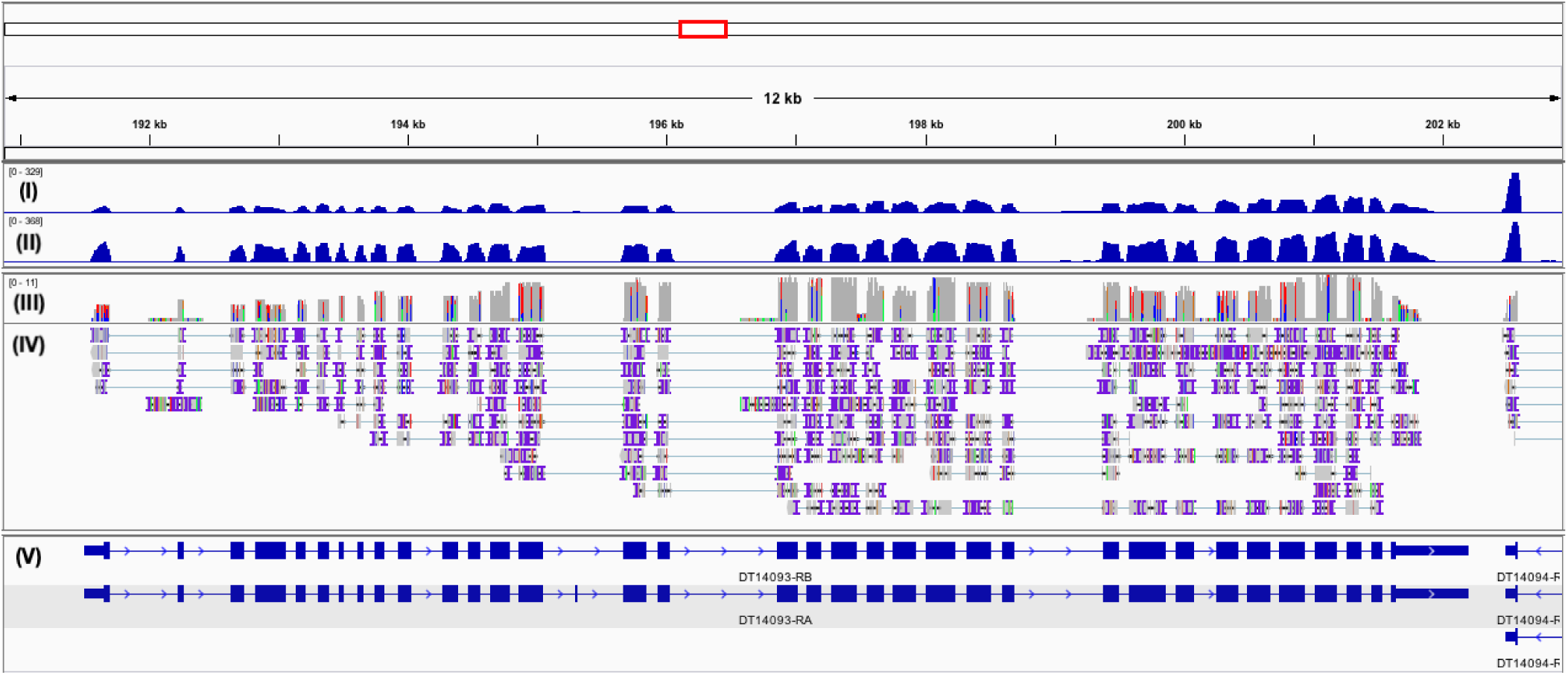
IGV screenshot of the *dnmt1* locus in the DT genome assembly, with short-read RNA coverage, mapped Nanopore RNA-seq reads and alternative splicing annotation. Tracks from top to bottom: (I) adult RNA-seq coverage, (II) 3 dpf RNA-seq coverage, (III) Nanopore RNA-seq coverage, (IV) Nanopore RNA-seq read mapping and (V) annotation with alternative splicing isoforms.

### DT and zebrafish intron size distributions

The predicted genome size of DT is around one half of the zebrafish reference genome (27). *Danionella dracula*, a close relative of DT, possesses a unique developmentally truncated morphology (28) and has a genome of a similar size (ENA accession number GCA_900490495.1). In order to validate our genome assembly, we set out to compare the compact genome of DT to the zebrafish reference genome.

Changes in the intron lengths have been shown to be a significant part of genomic truncations and expansions, such as a severe intron shortening in another miniature fish species, *Paedocypris* (29), or an intron expansion in zebrafish (30). We therefore compared the distribution of total intron sizes from the combined Ensembl/Havana zebrafish annotation (18) to the MAKER-produced DT annotation (Fig. 4a). We found that the DT intron size distribution is similar to other fish species investigated in ref. (29) which stands in stark contrast to the large tail of long introns in zebrafish. Median intron length values are in the range of the observed genome size difference (462 bp in DT as compared to 1,119 bp in zebrafish).

**Fig. 4.**
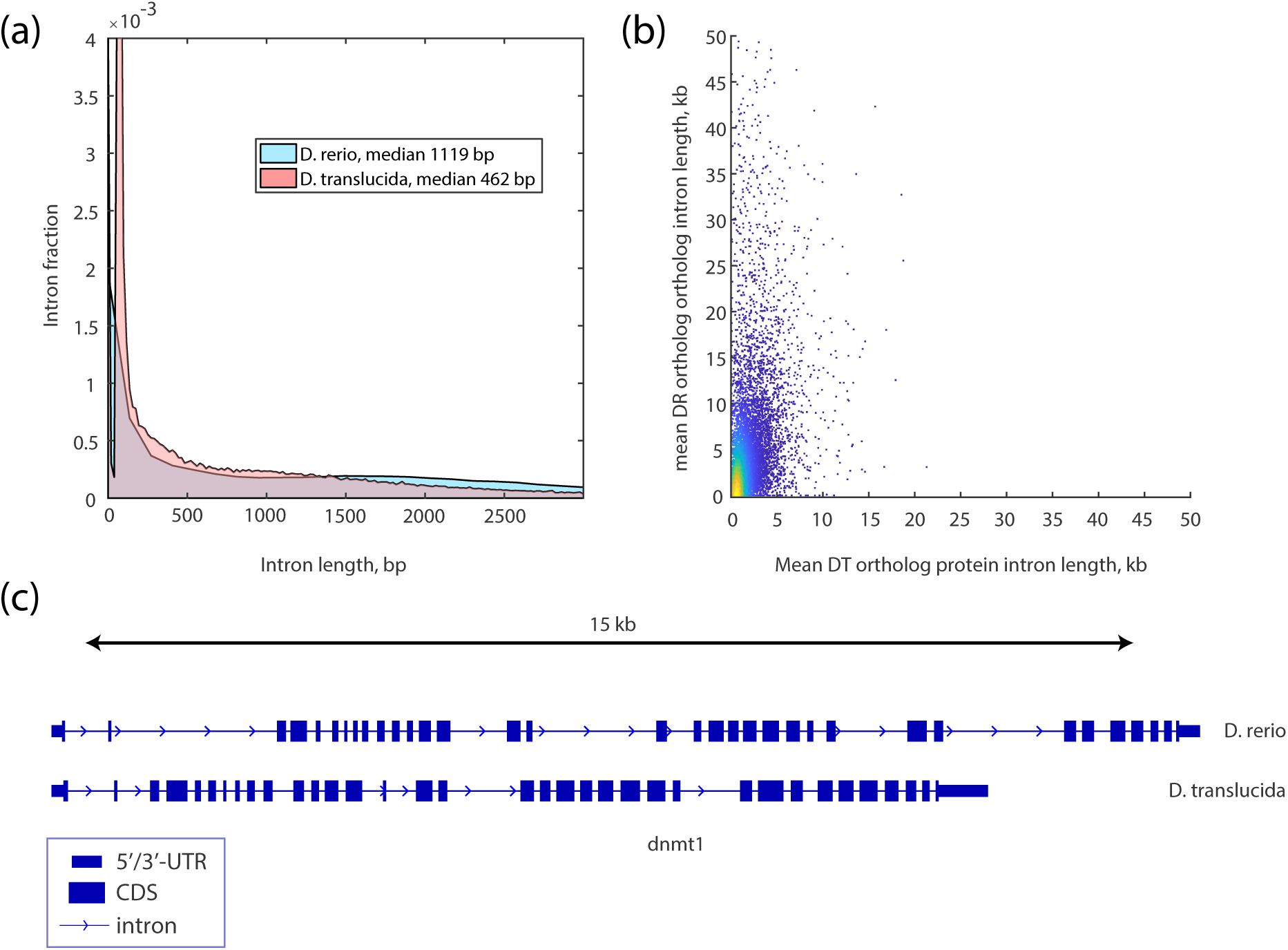
Intron size distribution in DT in comparison to zebrafish (DR). (a) Intron size distribution of all transcripts in DR and DT. (b) Intron size relationship for identified DR-DT orthologous proteins. (c) *dnmt1* ortholog locus in both fish. 5′/3′ UTR, untranslated regions; CDS, coding sequence.

To investigate the difference in intron sizes on the transcript level, we compared average intron sizes for orthologous protein-coding transcripts in DT and zebrafish. We have identified orthologs in DT and zebrafish protein databases with the help of the conditional reciprocal best BLAST hit algorithm (CRB-BLAST) (31). In total, we have identified 19,192 unique orthologous protein pairs. For 16,751 of those orthologs with complete protein-coding transcript exon annotation in both fish we calculated their respective average intron lengths (Fig. 4b). The distribution was again skewed towards long zebrafish introns in comparison to DT. As an example, Fig. 4c shows *dnmt1* locus for the zebrafish and DT orthologs.

## Conclusions

In this work we describe whole-genome sequencing, assembly and annotation of the *Danionella translucida* genome. Using deep-coverage short reads and low-coverage long Nanopore reads, we achieved a high level of assembly continuity and completeness. We have functionally annotated the assembly with both long- and short-read RNA-seq, which allowed us to quantify the intron size distribution in DT in comparison to zebrafish. We expect that this work will provide an important resource for the use of *Danionella translucida* in biomedical research.

## Data availability

The genome assembly and annotation files and data analysis codes will be made available on g-node.

## ACKNOWLEDGEMENTS

We would like to thank Jörg Henninger for helpful discussions and critical reading of this manuscript. This work was funded by the NeuroCure Cluster of Excellence (Exc. 257) to MS and BJ. BJ is a recipient of a Starting Grant by the European Research Council (ERC-2016-StG-714560) and the Alfried Krupp Prize for Young University Teachers, awarded by the Alfried Krupp von Bohlen und Halbach-Stiftung.

